# Low albumin status accompanies multi-layered immunosuppressive phenotypes in metastatic breast cancer patients

**DOI:** 10.1101/2023.09.05.556440

**Authors:** Yuki Nakamura, Mayuko Yoda, Yoshihiro Izumi, Yukie Kashima, Masatomo Takahashi, Kohta Nakatani, Takeshi Bamba, Chenfeng He, Riyo Konishi, Don Pietro Saldajeno, Alexis Vandenbon, Yutaka Suzuki, Masakazu Toi, Kosuke Kawaguchi, Shinpei Kawaoka

## Abstract

Low albumin status is prevalent in advanced cancer patients, but the pathophysiology associated with this anomaly remains largely unexplored. To address this, we aim to search correlations of albumin levels with the transcriptome against peripheral blood mononuclear cells and the plasma metabolome within the same patients having metastatic breast cancers. We confirm that metastatic breast cancer patients exhibit low albumin levels in varying degrees without prominent systemic inflammation. Our data demonstrate that low albumin levels correlate with transcriptome signatures indicative of “neutrophil activation and T-cell down-regulation,” an immunosuppressive phenotype. We also find that immunoregulatory metabolites, such as arginine, are reduced in plasma in an albumin-correlated manner, further corroborating systemic immunosuppression. These results are verified using a mouse model of breast cancer. We conclude that low albumin status in metastatic breast cancer patients accompanies immunosuppressive phenotypes, which is likely unfavorable for anti-cancer immunotherapy and thus can be a cause of unsuccessful treatment outcomes.

## Introduction

Managing quality of life and homeostasis in metastatic cancer patients is a crucial unmet clinical need^1, 2^. Advanced cancers cause abnormalities in various host organs and cells, disrupting host homeostasis^3, 4, 5, 6^. Such disruption includes rewired systemic metabolism and corruption in immunity and is clinically called cancer cachexia^1, 2, 3, 4, 6^. Disrupted homeostasis in patients generally leads to reduced quality of life and lowered efficacy of anti-cancer therapies including anti-cancer immunotherapy^7, 8^. Given the impacts of such cancer-induced disruption on a patient’s life and treatment strategies, it has been demanded to develop novel therapeutics that can mitigate cancer’s adverse effects on the host^1, 2^.

Our ability to accurately capture the patient’s status is currently limited. This is largely due to our insufficient understanding of the comprehensive picture of host pathophysiology in cancers. For example, it is known that blood albumin levels are low in patients with advanced cancers, which generally indicate poor nutritional condition and associate with poor prognosis^9, 10, 11, 12^. However, it remains incompletely understood what kind of pathophysiology is associated with albumin down-regulation in advanced cancer patients in detail. Deepening our understanding of such disrupted homeostasis (e.g., low albumin) is essential to find a way to manage the quality of life and treatment strategies in advanced cancer patients.

Comprehensive measurements of diverse genes and molecules (i.e., omics analysis) against host organs and cells in cancer-bearing conditions is a promising method to understand cancer’s adverse effects on the host. To date, a relatively non-biased approach led to the identification of altered metabolism in various metabolic organs, dysregulation in the immune system, and so on., as exemplified by but not limited to a series of previous reports^13, 14, 15, 16, 17, 18, 19^. Measuring molecules from different sources (e.g., blood and immune cells) is of particular significance because it would enable the thorough characterization of disrupted homeostasis. Yet, multi-omics studies that aim to understand host pathophysiology in cancers have been relatively rare, especially in patients.

In the current study, we aim to characterize the disrupted homeostasis of patients with metastatic breast cancers (stage IV). We recruited five healthy volunteers and ten patients with stage IV breast cancer. We performed gene expression analyses in peripheral blood mononuclear cells and metabolome analyses of the plasma in the same subjects. Correlating those multi-layered parameters indicated that low albumin levels are associated with the increase in neutrophils and decreases in CD8^+^ T cells, γδ T cells, and natural killer cells. This association did not accompany the elevation of C-reactive proteins (CRPs), the marker for systemic inflammation^20^. We also found that a set of immunoregulatory metabolites, such as arginine^21, 22^, correlates with low albumin levels. We validated these results using a mouse model of breast cancer. Collectively, the current study identifies multi-omics signatures associated with low albumin status, finding albumin-correlated systemic immunosuppression in metastatic breast cancer patients.

## Results

### Multi-layered characterization of metastatic breast cancer patients

We aimed to construct multi-layered omics datasets from metastatic breast cancer patients (stage IV) (Fig. 1a). We collected blood samples from five healthy volunteers and ten advanced breast cancer patients harboring visceral metastasis and performance status ≤ 2 (Table 1)^23^. To make the effects of circadian rhythm minimum, we collected the specimens at a similar time of day (Table 1: AM 11:11 ± 01:59). We isolated peripheral blood mononuclear cells (PBMC) and plasma, subjecting them to gene expression and metabolome analyses, respectively (Extended Data Table 1-4). We also recorded patients’ information, including the site of metastasis and ages. We integrated those rich datasets to understand the nature of host pathophysiology correlating with albumin levels in breast cancer patients.

**Figure 1:**
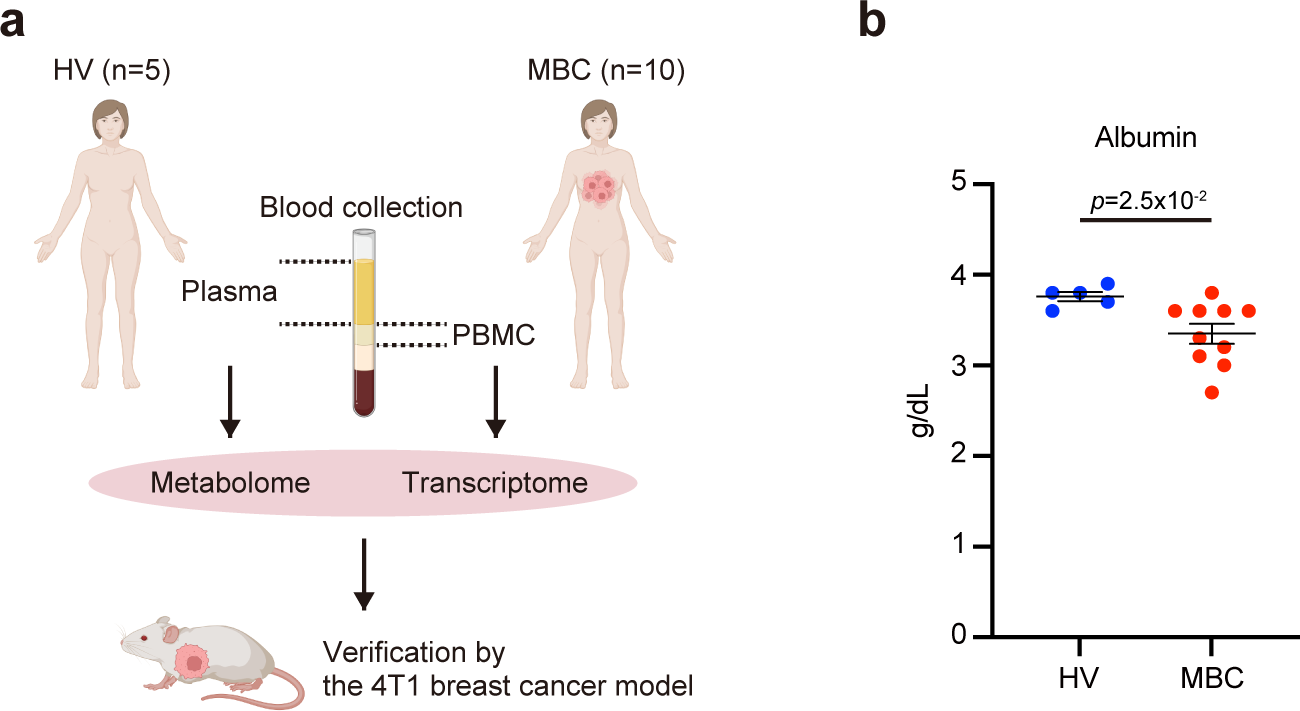
Metastatic breast cancers reduce plasma albumin levels. **a.** Design of this study. **b.** The plasma albumin levels of healthy volunteers (HV) and metastatic breast cancer patients (MBC). Data are presented as the mean ± SEM. The *p-*value is shown (non-paired, two-tailed Student *t*-test). *n* = 5 for HV, *n* = 10 for MBC. See also Extended Data Fig. 1a for the serum albumin levels from the same patients, and Extended Data Fig. 1b for the levels of C-reactive protein (CRP), a marker for systemic inflammation, from the same patients.

**Table 1.** Patient characteristics.

| ID |  | Time of blood collection | Age (years) | Sex | PS | Subtype |  | Metastatic site | No. of previous treatment regimens | Duration of treatment |
| --- | --- | --- | --- | --- | --- | --- | --- | --- | --- | --- |
|  |  |  |  |  |  | HR | HER2 |  |  |  |
| MBC | 1 | 11:00 | 71 | Female | 1 | + | - | Lung, liver, lymph node | 2 | 6 years |
|  | 2 | 16:00 | 77 | Female | 2 | + | - | Lung, liver, bone, lymph node | 10 | 9 years |
|  | 3 | 9:40 | 71 | Female | 1 | + | - | Lung, bone | 5 | 5 years |
|  | 4 | 13:20 | 70 | Female | 1 | + | - | Lung, liver, bone | 4 | 6 years |
|  | 5 | 10:30 | 39 | Female | 1 | + | - | Liver, bone, brain, ovary, adrenal glands, peritoneum | 0 | 2 years |
|  | 6 | 10:00 | 56 | Female | 1 | + | - | Lung, liver, bone | 0 | 6 months |
|  | 7 | 10:30 | 76 | Female | 1 | - | - | Lung, bone, lymph node | 1 | 1 year |
|  | 8 | 12:15 | 76 | Female | 2 | - | - | Lung, lymph node | 1 | 6 months |
|  | 9 | 11:00 | 51 | Female | 2 | - | + | Lung, liver, lymph node | 0 | 9 months |
|  | 10 | 11:00 | 64 | Female | 1 | + | + | Lung, Liver, bone, lymph node | 4 | 3 years |
| HV | 1 | 9:00 | 33 | Female | N.D. |  |  |  |  |  |
|  | 2 | 9:15 | 37 | Female |  |  |  |  |  |  |
|  | 3 | 9:30 | 28 | Female |  |  |  |  |  |  |
|  | 4 | 10:00 | 49 | Female |  |  |  |  |  |  |
|  | 5 | 14:45 | 36 | Female |  |  |  |  |  |  |
| Average |  | 11:11 (±1:59) | 55.6 (±17.7) |  |  |  |  |  |  |  |
| Average of MBC |  | 11:31 (±1:48) | 65.1 (±12.6) |  |  |  |  |  |  |  |
| Average of HV |  | 10:30 (±2:09) | 36.6 (±7.8) |  |  |  |  |  |  |  |
| *p*-value |  | 0.38 | 0.00052 |  |  |  |  |  |  |  |

Abbreviation: MBC, metastatic breast cancer patients
HV, healthy volunteers
PS, The Eastern Cooperative Oncology Group performance status
HR, hormone receptor

Our measurements for plasma albumin demonstrated that metastatic breast cancer patients had lower plasma albumin levels compared to healthy volunteers (Fig. 1b). The average in healthy volunteers was 3.8 ± 0.10 (g/dL), while that in breast cancer patients was 3.4 ± 0.33 (*p* = 0.025). We validated that these patients had similar serum albumin levels, lending credence to our measurements (Extended Data Fig. 1a). It was of note that the standard deviation in breast cancer patients was larger than healthy volunteers, indicating that breast cancer patients were more heterogenous regarding albumin levels than healthy volunteers. This result strongly suggested that host pathophysiology (e.g., nutritional status) significantly differed among metastatic breast cancer patients even though they were grouped as stage IV, demanding the need to stratify those heterogenous patients. It should be additionally noted that only three of ten patients had > 0.3 mg/dL C-reactive protein (CRP) levels (Extended Data Fig. 1b), indicating that systemic inflammation was not evident in this patient cohort. Based on these, we decided to search for multi-omics signatures associated with low albumin status.

### Identification of transcriptome signatures correlating with low albumin status

To find PBMC transcriptome signatures correlating with albumin levels, we utilized weighted gene co-expression network analysis (WGCNA)^24^. WGCNA allows us to find modules that consist of genes with similar expression patterns. We can also correlate a particular parameter (e.g., albumin levels) and co-expression modules. In our analyses, we wanted to focus on annotated protein-coding genes that were potentially differentially expressed between healthy volunteers and metastatic breast cancer patients. For this purpose, we selected genes exhibiting more than 1.5-fold differences, resulting in 2,117 genes for our WGCNA analyses (Extended Data Fig. 2a, b). WGCNA identified seven co-expression modules (Fig. 2a). We included age as a parameter because we were aware of the age difference between the two groups (Table 1). Among the modules, we found a strong negative correlation between module-3 and albumin scores (Fig. 2a; *r* = −0.74 and *p* = 0.002). On the other hand, this module did not correlate with age (*r* = 0.089 and *p* = 0.8). Module-3 contained 155 genes (Fig. 2b). When we plotted these 155 genes according to their correlation to albumin levels, we found that 146 genes in this module had negative correlations with albumin levels and were up-regulated in metastatic breast cancer patients (Fig. 2b). On the other hand, 9 genes showed positive correlations with albumin.

**Figure 2:**
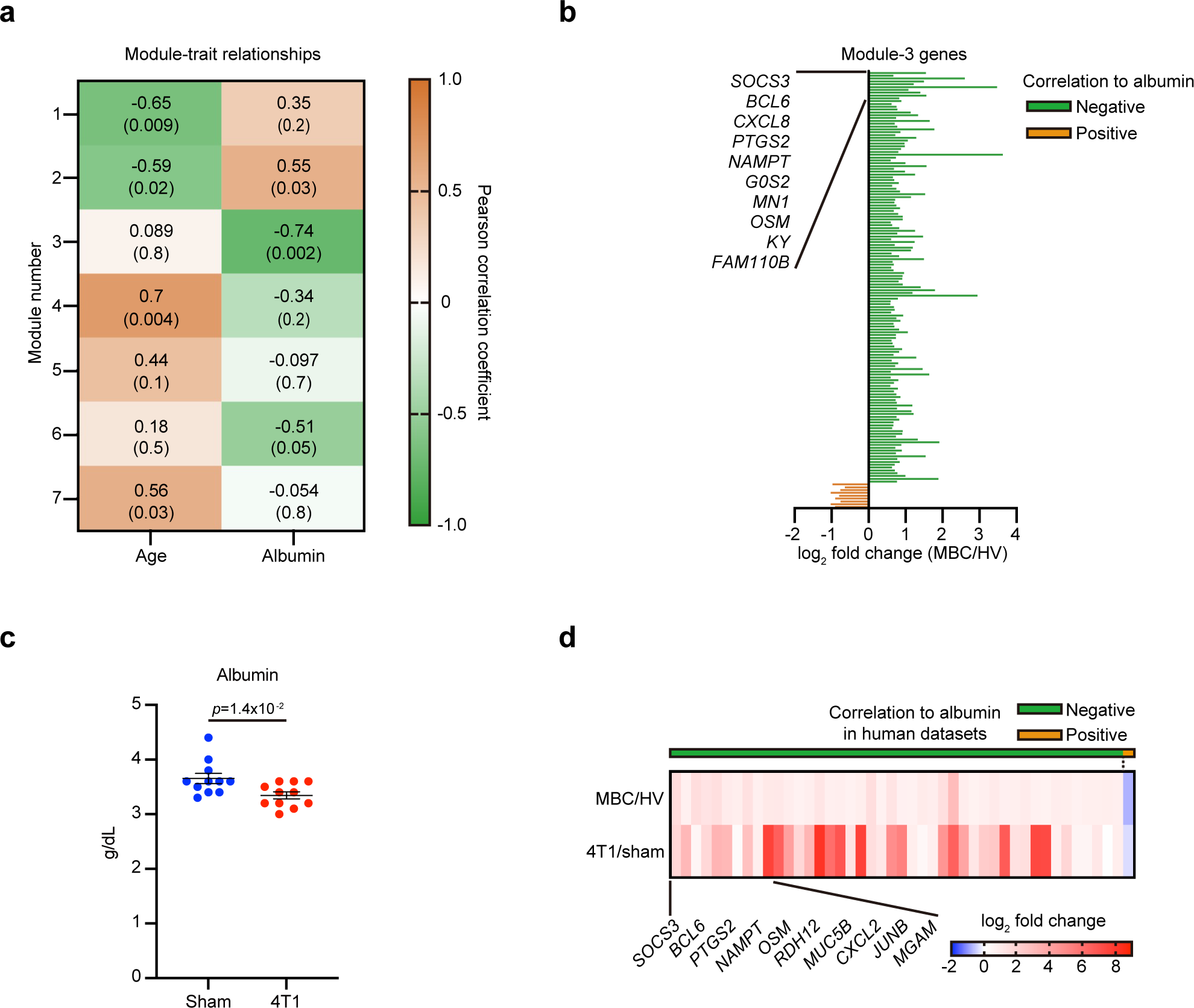
Identification of gene modules correlating with albumin levels. **a.** Heatmap showing the Pearson correlation coefficients between the clinical characteristics (age and albumin levels) and the module eigengenes generated by WGCNA. In each column, the upper number indicates the Pearson correlation coefficient and the lower number in parentheses indicates a *p*-value calculated by WGCNA. See also Extended Data Fig. 2a, b for scale independence and mean connectivity of this analysis. **b.** Bar plot showing log_2_ fold changes of module-3 genes between metastatic breast cancer patients (MBC) and healthy volunteers (HV). Genes showing negative correlations to albumin levels are shown in green. Genes showing positive correlations to albumin levels are shown in orange. Genes are ordered according to their correlation to albumin levels. The names of the top10 genes most strongly negatively correlating with albumin levels are indicated. **c.** The plasma albumin levels of sham-operated and 4T1 breast cancer-bearing mice measured 14 days after transplantation. Data are presented as the mean ± SEM. The *p-*value is shown (non-paired, two-tailed Student *t*-test). *n* = 11 for sham-operated mice, *n* = 11 for 4T1-bearing mice. **d.** Heatmap of module-3 genes showing the same direction of changes in the 4T1 breast cancer model compared to the human datasets. Genes are ordered according to their correlation to albumin levels in humans. The names of the top10 genes most strongly negatively correlating with albumin levels in humans are indicated. See also Extended Data Fig. 2c, d for the mouse data.

We then investigated whether module-3 genes were also differentially expressed in a mouse model of breast cancer (4T1)^25^. This analysis was to validate our human transcriptome datasets and to consider the effects of e.g., chemotherapy in human datasets. Due to various different treatment conditions and health statuses in patients, it is sometimes challenging to attribute identified host changes to cancers. Mouse cancer models are simpler than patients and do not have this problem, helping identify changes owing to cancers in patients.

We found that plasma albumin levels in BALB/c mice were severely reduced on day 14 post-transplantation of 4T1 breast cancer cells (Fig. 2c). We performed transcriptome analyses against PBMC of sham-operated and 4T1-bearing mice, obtaining differentially expressed genes (Extended Data Fig. 2c). We then sought module-3 genes commonly differentially expressed in humans and mice, obtaining 45 genes (Fig. 2d). Most of these mouse genes were up-regulated in 4T1-bearing mice in which albumin was down-regulated (Extended Data Fig. 2d), as was the case for humans (Fig. 2b). Successful identification of PBMC genes differentially expressed under low albumin conditions in humans and mice allowed us confidently characterize host pathophysiology in human breast cancer patients.

### Low albumin status accompanies neutrophil activation

We further explored module-3 genes to understand what this module represented in detail. To this end, we examined correlations between albumin levels and module-3 genes individually (Fig. 3a-c). We also wanted to know which immune cell type expressed these genes (Fig. 3d-f). For this purpose, we exploited PBMC single-cell RNA-seq (scRNA-seq) datasets obtained from Japanese healthy female volunteers that were previously published^26^. This analysis identified 18 cell types (Extended Data Fig. 3a). As exemplified in Fig. 3a-c, representative module-3 genes (*SOCS3, BCL6*, and *PTGS2*; Fig. 2d) showed strong negative correlations with albumin levels. These genes showed different expression patterns in PBMC, but all were expressed in neutrophils that were defined by the restricted expression of *S100A8* (also known as *MRP8*) to the cluster previously annotated as classical monocytes (Fig. 3d-f and Extended Data Fig. 3a, b)^26, 27^. Of note, the expression of *PTGS2*, an inflammation-related gene encoding cyclooxygenase proteins^28^, appeared relatively specific to neutrophils (Fig. 3f and Extended Data Table 5). *BCL6* also had a clear relation to neutrophils: a recent study revealed that *Bcl6* in neutrophils promotes inflammation by extending neutrophil survival, worsening host pathophysiology in a mouse model of virus infection^29^. The expression of *BCL6* was biased toward neutrophils as well (Fig.3e). These results led to the idea that low albumin status might correlate with neutrophils.

**Figure 3:**
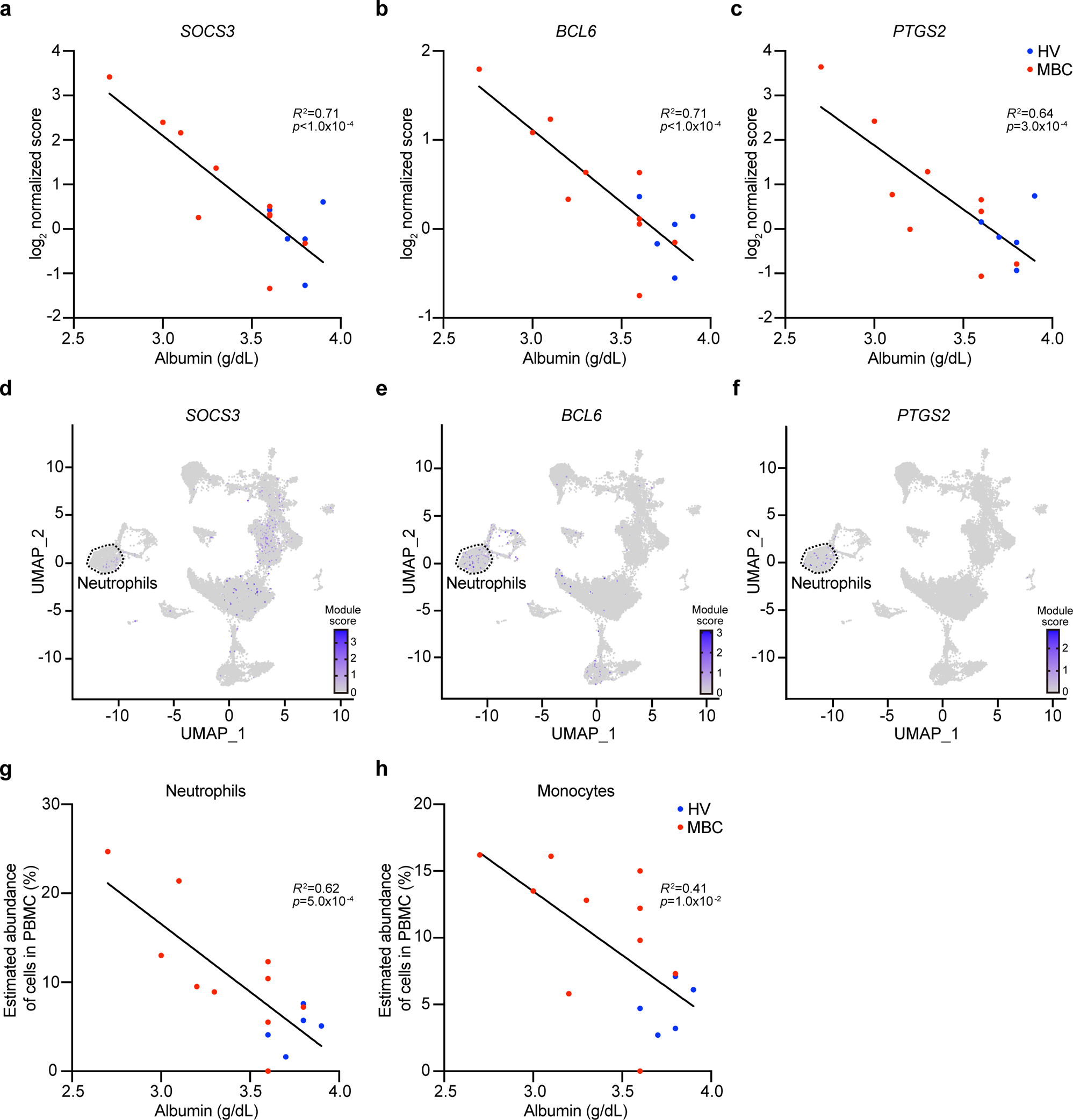
Low albumin status accompanies a signature of neutrophil activation. **a.** Correlation of plasma albumin levels and log_2_ normalized scores of *SOCS3* in PBMC. **b.** Correlation of plasma albumin levels and log_2_ normalized scores of *BCL6* in PBMC. **c.** Correlation of plasma albumin levels and log_2_ normalized scores of *PTGS2* in PBMC. **d.** UMAP plot for *SOCS3*. **e.** UMAP plot for *BCL6*. **f.** UMAP plot for *PTGS2*. **g.** Estimated abundance of neutrophils in PBMC calculated by ImmuCell-AI and its correlation to plasma albumin levels. **h.** Estimated abundance of monocytes in PBMC calculated by ImmuCell-AI and its correlation to plasma albumin levels. See also Extended Data Fig. 3e, g-i for the results from other immune cell types. **a-c.** Scores are normalized to the average scores of five healthy volunteers. **a-c, g, h.** The Pearson correlation coefficients and *p*-values obtained by simple regression analysis (GraphPad Prism) are shown. **d-f.** The data are retrieved from previously published single-cell RNA-seq (scRNA-seq) datasets from PBMC of Japanese healthy volunteers (*n* = 3 (females))^26^. A cluster corresponding to neutrophils is highlighted. See also Extended Data Fig. 3b for neutrophil identification in this UMAP plot.

To further examine the immune cell status in metastatic breast cancer patients, we analyzed the proportion of various immune cell types using ImmuCell-AI^30, 31^. We calculated the estimated abundance of immune cells within PBMC for neutrophils, monocytes, dendritic cells, macrophages, natural killer cells, natural killer T cells, CD4^+^ T cells, CD8^+^ T cells, γδ T cells, and B cells. Consistent with our gene expression analyses (Fig. 3a-f), neutrophils in PBMC were increased in metastatic breast cancer patients in a manner correlated with albumin levels (Fig. 3g). The increase in neutrophils in the cancer-bearing condition was validated using the 4T1 breast cancer model (Extended Data Fig. 3c). In addition, we found that monocytes defined by ImmuCell-AI were negatively correlated with albumin levels (Fig. 3h and Extended Data Fig. 3d). Dendritic cells also showed a negative correlation with albumin (Extended Data Fig. 3e). Module-3 genes were indeed expressed in dendritic cells based on the human protein atlas (Extended Data Table 5), but this cell type was reduced rather than increased in the 4T1 model (Extended Data Fig. 3f). Macrophages, natural killer T cells, and CD4^+^ T cells did not show strong correlations with albumin levels (Extended Data Fig. 3g-i). Collectively, together with the elevations of *SOCS3*, *PTGS2*, and *BCL6*, these results suggest that module-3 genes represented “elevated cellular inflammatory and anti-apoptosis signals in increasing neutrophils.” Of note, as described earlier, patients having these signatures did not show strong systemic inflammation (i.e., high CRP levels; Extended Data Fig. 1b)^28, 29, 32^.

### Low albumin status accompanies deficiencies in CD8^+^ T cells and γδ T cells

We next examined module-2 genes that were positively correlated with albumin and negatively correlated with age (Fig. 2a). The involvement of age was considered by excluding individual genes showing strong correlations with age (|*r*| > 0.5) and by examining the 4T1 datasets that were free from age differences. We found that most of the resultantly selected 154 module-2 genes were down-regulated in PBMC of metastatic breast cancer patients and 4T1-bearing mice (Fig. 4a). In this analysis, *CD8A*, the marker for CD8^+^ T cells^33^, got our attention. Our data demonstrated that representative module-2 genes, including *CD8A*, were positively correlated with albumin in line with the WGCNA result (Fig. 4b-d and Fig. 2a). scRNA-seq datasets confirmed enrichment of these module-2 genes in CD8^+^ T cells in healthy volunteers (Fig. 4e-g). *KLRG1* and *CRTAM* were suggestive of exhaustion in CD8^+^ T cells according to previous studies^34, 35^. ImmuCell-AI revealed that CD8^+^ T cells in PBMC were reduced in a manner associated with albumin levels in metastatic breast cancer patients (Fig. 4h). CD8^+^ T cells were decreased in PBMC of 4T1-bearing mice as well (Extended Data Fig. 4a). These results indicated that module-2 genes represented “down-regulation and exhaustion of CD8^+^ T cells in PBMC” in a manner correlated with albumin. We suggest that these changes represent systemic immunosuppression in metastatic breast cancer patients, which will be discussed in detail in the following sections.

**Figure 4:**
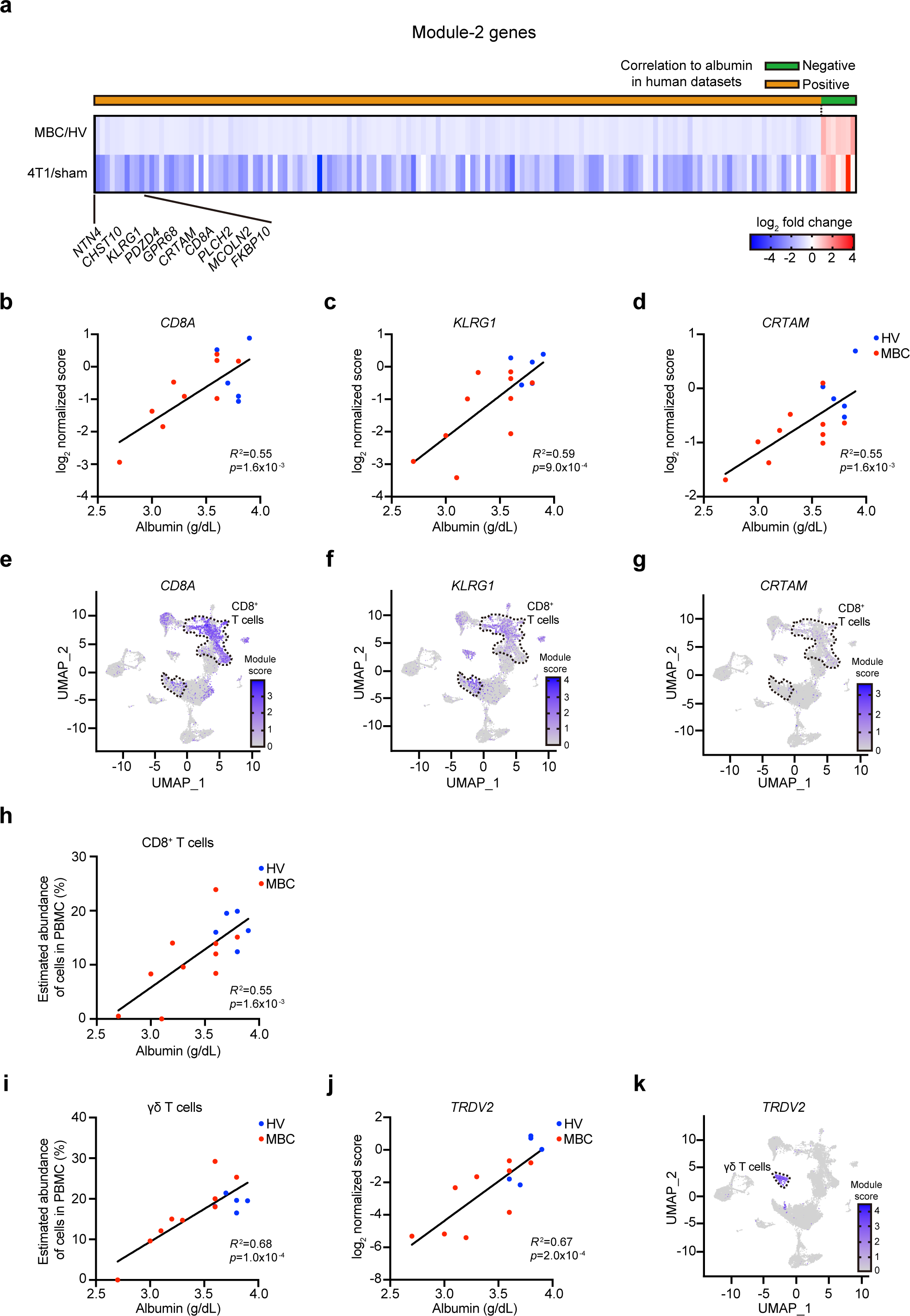
Low albumin status accompanies a signature of T cell suppression. **a.** Heatmap of select module-2 genes showing the same direction of changes in the 4T1 breast cancer model compared to the human datasets. Genes are ordered according to their correlation to albumin levels in humans. The names of the top10 genes most strongly positively correlating with albumin levels in humans are indicated. **b.** Correlation of plasma albumin levels and log_2_ normalized scores of *CD8A* in PBMC. **c.** Correlation of plasma albumin levels and log_2_ normalized scores of *KLRG1* in PBMC. **d.** Correlation of plasma albumin levels and log_2_ normalized scores of *CRTAM* in PBMC. **e.** UMAP plot for *CD8A*. **f.** UMAP plot for *KLRG1*. **g.** UMAP plot for *CRTAM* **h.** Estimated abundance of CD8^+^ T cells in PBMC calculated by ImmuCell-AI and its correlation to plasma albumin levels. **i.** Estimated abundance of γδ T cells in PBMC calculated by ImmuCell-AI and its correlation to plasma albumin levels. See also Extended Data Fig. 4b for the data related to natural killer cells. See also Extended Data Fig. 4a, c, d for the mouse data for CD8^+^ T cells, γδ T cells, and natural killer cells, respectively. **j.** Correlation of plasma albumin levels and log_2_ normalized scores of *TRDV2* in PBMC. **k.** UMAP plot for *TRDV2*. **b-d, j.** Scores are normalized to the average scores of five healthy volunteers. **b-d, h, i, j.** The Pearson correlation coefficients and *p*-values obtained by simple regression analysis (GraphPad Prism) are shown. **e-g, k.** The data are retrieved from previously published single-cell RNA-seq (scRNA-seq) datasets from PBMC of Japanese healthy volunteers (*n* = 3 (females))^26^. Clusters corresponding to CD8^+^ T cells (**e-g**) and γδ T cells (**k**) are highlighted.

ImmuCell-AI analyses identified two more cell types showing positive correlations with albumin: γδ T cells and natural killer cells (Fig. 4i and Extended Data Fig. 4b). Markers for γδ T cells such as *TRDV2*^36^ exhibited strong positive correlations with albumin levels (Fig. 4j, k). The reduction in γδ T cells was observed in the 4T1 model as well (Extended Data Fig. 4c). Natural killer cells were also decreased in the 4T1 model (Extended Data Fig. 4d), and a set of module-2 genes were indeed expressed in this cell type (Extended Data Table 6). Although the degree of correlation between albumin and natural killer cells was lesser compared to CD8^+^ T cells and γδ T cells, these results suggested that natural killer cells were likely affected by cancers in an albumin-correlated manner. Based on these results, we concluded that module-2 genes represented alterations in γδ T cells and natural killer cells in addition to alterations in CD8^+^ T cells.

It was an intriguing contrast that low albumin status was negatively correlated with neutrophils but positively correlated with CD8^+^ T cells, γδ T cells, and natural killer cells. We interpreted this as an immune cell imbalance associated with albumin in metastatic breast cancer patients. To further explore such imbalance in patients, we calculated ratios among these cell types within PBMC. As shown in Fig. 5, we found that the ratio between neutrophils and CD8^+^ T cells, neutrophils and γδ T cells, and neutrophils and natural killer cells was strongly negatively correlated with albumin (Fig. 5a-c). To validate these results, we analyzed data from patients’ charts, calculating the neutrophil-to-lymphocyte ratio in whole blood, a parameter known as NLR^37^. This analysis confirmed a negative correlation between NLR and albumin (Extended Data Fig. 5). Although the ratios within PBMC and NLR in whole blood are different biological parameters, their correlations with albumin levels were consistent. In summary, our data revealed correlations between albumin levels and immune cell imbalance in PBMC, further implicating systemic immunosuppression in metastatic breast cancer patients.

**Figure 5:**
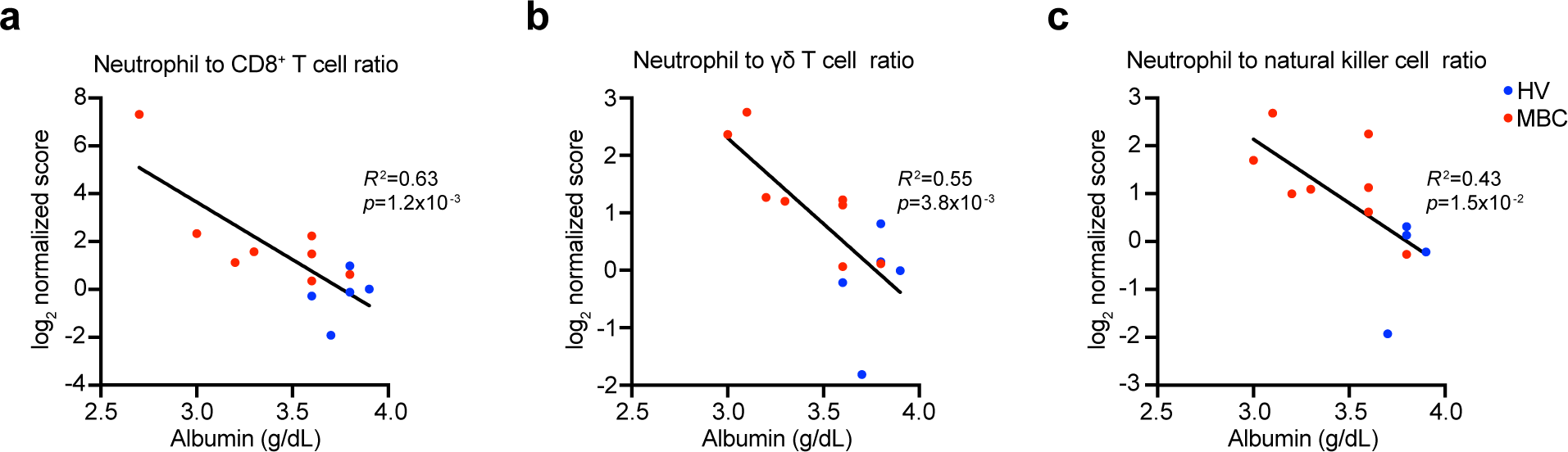
Low albumin status accompanies an immune cell imbalance. **a.** Correlation between plasma albumin levels and neutrophil-to-CD8^+^ T cell ratio in PBMC. **b.** Correlation between plasma albumin levels and neutrophil-to-γδ T cell ratio in PBMC. **c.** Correlation between plasma albumin levels and neutrophil-to-natural killer cell ratio in PBMC. **a-c.** The Pearson correlation coefficients and *p*-values obtained by simple regression analysis (GraphPad Prism) are shown. Note that data points showing uncalculatable infinite scores (i.e., denominator is 0) are not shown in the figure. See also Extended Data Fig. 5 for neutrophil-to-lymphocyte ratio in whole blood (i.e., NLR) calculated from patients’ charts.

### Low albumin status accompanies immunosuppressive metabolic phenotypes

We next wanted to address if metabolic changes in plasma could further support our notion on albumin-correlated immune cell imbalance and immunosuppression in metastatic breast cancer patients. For this purpose, we performed metabolome analyses against plasma from metastatic breast cancer patients and 4T1-bearing mice (Fig. 6 and Extended Data Table 4). We first analyzed 126 metabolites in metastatic breast cancer patients to find metabolites correlating with albumin levels in patients (Fig. 6a). This analysis identified 10 metabolites showing positive or negative correlations with albumin levels when |*r*| > 0.5 was set as a threshold. We then tried to find overlaps between human datasets and 4T1 mouse datasets. Among 10 metabolites listed from the human datasets, 7 metabolites exhibited the same direction of changes in the plasma of 4T1-bearing mice (Fig. 6b). These metabolites included aspartate, glutamic acid, hippuric acid, ketoleucine, methionine, arginine, and azelaic acid. Although it was difficult to confidently reveal the origin of cells and organs of these changes, we found a couple of informative metabolic changes that could be correlated to the immunosuppressive phenotypes in PBMC (Fig. 3-5).

**Figure 6:**
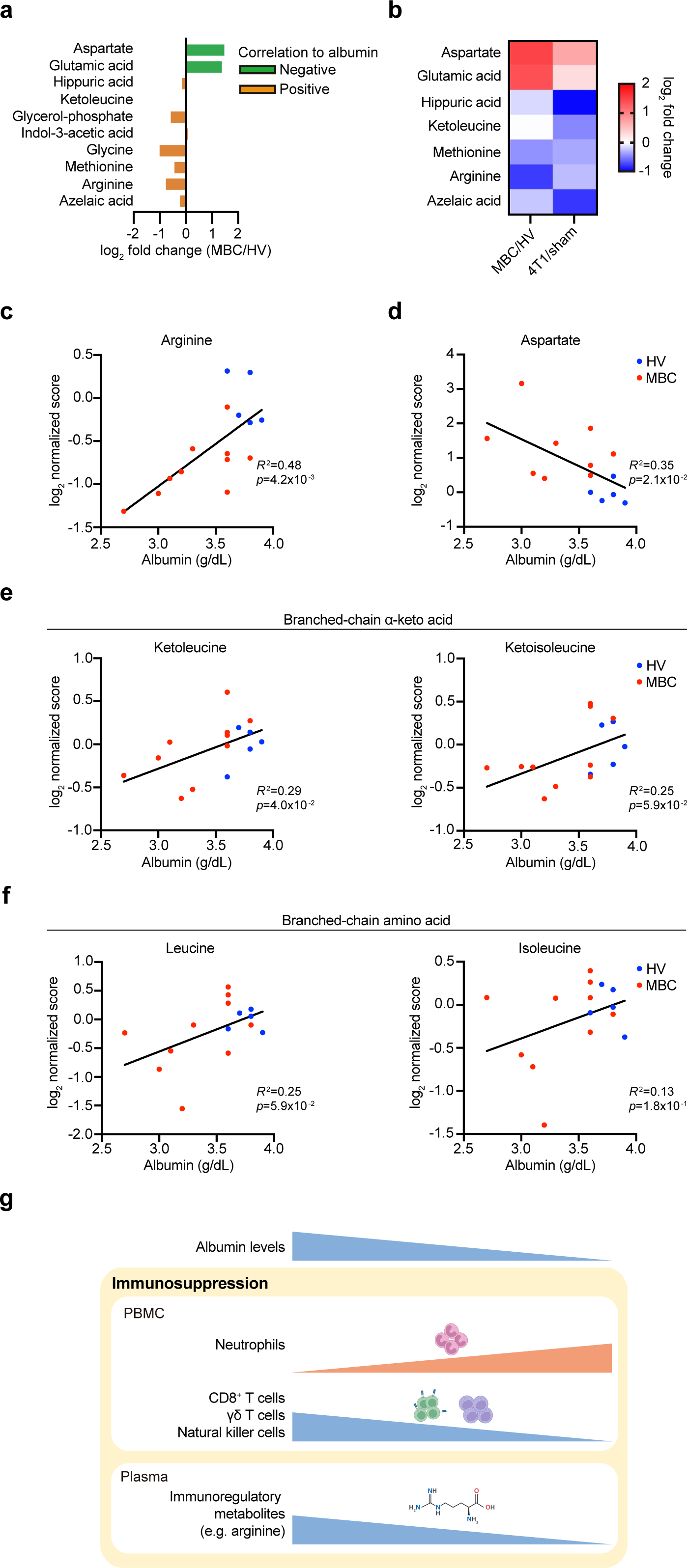
Low albumin status accompanies immunosuppressive metabolic phenotypes. **a.** Bar plot showing log_2_ fold changes of plasma metabolites between metastatic breast cancer patients (MBC) and healthy volunteers (HV). Metabolites showing negative correlations to albumin levels are shown in green (*r* < −0.5). Metabolites showing positive correlations to albumin levels are shown in orange (*r* > 0.5). Metabolites are ordered according to their correlation to albumin levels. **b.** Heatmap of metabolites shown in Fig. 6a that exhibit the same direction of changes between humans and mice. Metabolites are ordered according to their correlation to albumin levels in humans. **c.** Correlation of plasma albumin levels and log_2_ normalized scores of plasma arginine. **d.** Correlation of plasma albumin levels and log_2_ normalized scores of plasma aspartate. **e.** Correlation of plasma albumin levels and log_2_ normalized scores of plasma ketoleucine and ketoisoleucine.f. Correlation of plasma albumin levels and log_2_ normalized scores of plasma leucine and isoleucine. **g.** Summary of this study. **c-f.** Scores are normalized to the average scores of five healthy volunteers. The Pearson correlation coefficients and *p*-values obtained by simple regression analysis (GraphPad Prism) are shown.

Arginine showed a clear positive correlation with albumin levels and was reduced in cancer-bearing conditions (Fig. 6a-c). Arginine is a critical regulator of T cell-mediated immune responses, including anti-cancer immunity, and its depletion causes defects in T cells^21, 22, 38, 39^. Accordingly, arginine consumption by other cell types e.g., neutrophils can be a non-cell autonomous T-cell suppression mechanism^40^. It is known that activated neutrophils in patients with sepsis consume arginine, consequently limiting arginine availability for T cells^40^. Based on these, systemic reduction in arginine accompanied by neutrophil activation and the reduction in CD8^+^ T cells could be regarded as an interrelated immunosuppressive phenotype. Moreover, arginine has a different face as a member of the urea cycle in the liver^13,41^. Aspartate also contributes to the urea cycle^13, 41^ and was negatively correlated with albumin levels (Fig. 6d). Given previous studies reporting urea cycle dysfunction in patients and mouse models^13, 41^, we suggest that our plasma metabolome might have captured the previously reported abnormalities in the liver metabolism under the cancer-bearing condition. This assumption is in line with the fact that metastatic breast cancer patients had low albumin levels, suggestive of liver abnormalities.

Our data also demonstrated the alterations in the metabolism of branched-chain amino acids (BCAAs). BCAAs include valine, leucine, and isoleucine which are essential for organisms^42, 43^. It has been shown that BCAAs activate T cell types via mTOR signaling and glucose metabolism^44, 45, 46^. BCAAs are metabolized into branched-chain α-keto acids (BCKAs) in e.g., the livers and muscles and play roles in immunity as well as BCAAs^42, 43, 47^. We found that ketoleucine, a BCKA, was positively correlated with albumin levels and thus tended to be decreased in metastatic breast cancer patients (Fig. 6a, b, e). Ketoisoleucine exhibited a similar trend (Fig. 6e: *p* = 0.059). In contrast, corresponding BCAAs leucine and isoleucine did not show strong correlations with albumin levels (Fig. 6f). Yet, there was a weak tendency for these BCAAs to be reduced in some patients. These results suggested an altered systemic metabolism of BCAAs and BCKAs, further supporting that albumin-associated metabolome signatures in metastatic breast cancer patients implicate an immunosuppressive systemic environment in patients.

Our analyses captured changes in other metabolic pathways. For example, the tryptophan-kynurenine pathway is possibly affected by breast cancers (Extended Data Fig. 6b, c). We found that plasma tryptophan levels were decreased while plasma kynurenine levels tended to be increased in metastatic breast cancer patients (Extended Data Fig. 6b, c). These two amino acids have been implicated in the immune system^48^, as well as arginine and BCKAs. Although we require further functional studies to explore the role of these metabolic alterations in the immunosuppressive phenotypes, our datasets will be a useful resource for research on cancer-dependent host alterations in humans.

## Discussion

### Understanding low albumin status in metastatic breast cancer patients

Low albumin status is a prevalent phenotype in various diseases^9, 10, 11, 12^. Indeed, this is one of the most unfavorable symptoms, suggestive of low nutrition status, observed in patients with advanced cancers^9, 10, 11, 12^. In breast cancers, low albumin status is a prognostic factor^9, 10, 11, 12^. Despite its prevalence, what low albumin status tells us in detail in patients has been largely unexplored. In the current study, we aimed to tackle this issue with the aid of relatively unbiased measurements of gene expression in peripheral blood mononuclear cells and plasma metabolome.

The essence of our approach is to search correlations between albumin levels and multi-omics signatures rather than comparing grouped healthy volunteers and metastatic breast cancer patients (Fig. 1-2). Metastatic breast cancers are categorized as stage IV. However, the site of metastasis and treatment options differ among patients (Table 1). Furthermore, “the disease time course (where a patient is in a time course toward irreversibly disrupted homeostasis (i.e., death))” must be different among patients. Basically, we cannot control which disease time course we investigate in patients, which is a fundamental problem of clinical sample analyses. We tried to overcome this problem by correlating albumin levels, quantitative variables in patients, to multi-omics signatures obtained in a relatively unbiased manner. The small sample size is a limitation of our study, but our integrative analyses with the 4T1 murine breast cancer model helped the identification of signatures affected by cancers in patients.

Due to the prevalence of low albumin status in diseases^49^, our observations could be generalizable to other cancer types and diseases, such as sepsis and COVID-19. Our datasets thus will be useful in the research of albumin-correlated immunosuppression in other diseases beyond cancer, promoting personalized characterization of pathophysiology in diseases and treatment choices in clinics.

### Immunosuppressive phenotypes in metastatic breast cancer patients

Based on our data, we consider multi-omics signatures that we found correlated with albumin levels as “immunosuppressive phenotypes.” This interpretation relies on our observation that CD8^+^ T cells were down-regulated in patients and mice (Fig. 4-5). It is an interesting coincidence to find the reduction in arginine, which is the critical regulator of T cell activity^21, 22, 38, 39^. Previous studies revealed that arginine availability is crucial for T cells to exert their activity in culture and in vivo^38, 39^. Due to this arginine dependency, the consumption of arginine by other cell types can be a mechanism of T-cell suppression. In fact, arginine consumption by activated neutrophils is a known non-cell autonomous mechanism to suppress T cells^40^. The increase in neutrophils in PBMC appeared to match this model (Fig. 3). Such a cell-cell competition for arginine can be a feedback system to end immune responses but can also be an immunosuppression target by cancers to evade the host immunity.

Our previous study and others found that cancer transplantation in mice leads to a massive accumulation of neutrophils and suggested that these neutrophils can be classified as myeloid-derived suppressor cells (MDSCs)^14, 50^. Despite non-definitive, myeloid-derived suppressor cells are sometimes described as pathologically activated neutrophils^50^, which might correspond to the neutrophil activation phenotypes in our datasets. Furthermore, neutrophils detected in PBMC (also known as low-density neutrophils) in mouse cancer models show immunosuppressive activity^51^. These previous studies led to an assumption that the increased neutrophils in metastatic breast cancer patients might have a cellular function as MDSCs to suppress T-cell-mediated adaptive immunity in addition to arginine-depletion-dependent indirect suppression of T cells. These altogether give rise to an interpretation that the combination of neutrophil activation and T-cell down-regulation, which is accompanied by the reductions in immunoregulatory metabolites, suggests systemic immunosuppression in metastatic breast cancer patients (Fig. 6g). It will be critical to uncover the hierarchy among neutrophil activation, T cell suppression, and the reduction of immunoregulatory metabolites in plasma.

The previous studies demonstrated that prognosis is negatively correlated with the whole blood neutrophil-to-lymphocyte ratio (NLR) in breast cancer patients^37^. In addition, pre-treatment NLR is associated with the efficacy of anti-cancer immunotherapy in various cancer types, including breast cancer^52^. These studies supported our notion that the transcriptomically defined imbalance between neutrophils and lymphocytes in PMBC (Fig. 5) is a sign of worsened patients’ homeostasis, which is possibly not suitable for anti-cancer immunotherapy. Moreover, our study is important in that we found such immune imbalance in an unbiased manner rather than the hypothetical approach: neutrophils and CD8^+^ T cells are the most strongly correlated signatures with albumin levels. Causal relationships between low albumin status and the immunosuppressive phenotypes await further studies.

We do not exclude the possibility of the involvement of other non-immune cell types, such as hepatocytes and muscle cells, in the immunosuppressive phenotypes. Hepatocytes are the major site of albumin production^14^. Thus, low albumin status most likely indicates liver abnormalities. We previously showed that arginine accumulated in the livers of 4T1 breast cancer-bearing mice, which was related to cancer-dependent suppression of the urea cycle^13, 41^. Such liver deficiencies may explain the reduced availability of arginine in plasma, indirectly limiting arginine for T cells. The muscle is a source of branched-chain amino acids and branched-chain α-keto acids^42, 43, 47^. Therefore, low concentrations of these metabolites in plasma can be attributed to potential abnormalities in the muscle. Deciphering such inter-organ mechanism to explain immunosuppression in patients require further studies in the future.

### Local-to-systemic immunosuppression in advanced breast cancer patients

Our ongoing studies, including this study, indicate that breast cancers ingeniously suppress the host immunity during its progression. We recently reported that breast cancer metastases to the lymph nodes reduce the number of CD169^+^ lymph node sinus macrophages^53^. This macrophage is an initiator of anti-cancer immunity as they phagocytose cancer-derived antigens and present them to CD8^+^ T cells^54^. We regard the elimination of CD169^+^ macrophages as a mechanism of immunosuppression by breast cancers, which we found is prevalent in all breast cancer subtypes^53^. Our current study revealed the down-regulation of another crucial player of anti-cancer immunity, CD8^+^ T cells, in PBMC in metastatic breast cancer patients. Taken together, we suggest that advanced breast cancers can cause severe suppression of anti-cancer immunity locally and systemically. Moreover, γδ T cells and natural killer cells play roles in anti-cancer immunity^55, 56^. It is thus plausible that the down-regulation of these cell types also results in the suppression of anti-cancer immunity^5^.

As discussed above, defects in CD169^+^ macrophages and CD8^+^ T cells are considered corruptive in anti-cancer immunotherapy e.g., using anti-PD1 antibodies. Our data thus indicate that metastatic breast cancer patients with low albumin status have a systemic immune environment unfavorable for anti-cancer immunotherapy, which may explain unsuccessful cases of anti-cancer immunotherapy in breast cancer patients. Hence, understanding local and systemic mechanisms behind immunosuppression and developing novel therapeutics to combat such immunosuppression will help improve patients’ homeostasis and contribute to anti-cancer immunotherapy.

In summary, our study uncovered the albumin-associated immunosuppressive phenotypes in metastatic breast cancer patients using PBMC transcriptome and plasma metabolome, shedding light on the critical aspect of multi-layered host pathophysiology in breast cancers.

## Methods

### Human samples

All samples from healthy volunteers and metastatic breast cancer patients were collected at the Department of Breast Surgery, Kyoto University Hospital. Plasma and peripheral blood mononuclear cells (PBMC) were collected from patients having metastatic breast cancer during treatments. Written informed consent was given by all participants before collection. All study protocols were approved by the Institutional Ethical Committee (G0424-18) (Kyoto University Graduate School and Faculty of Medicine) and adhered to the provisions of the Declaration of Helsinki in 1995. The clinical characteristics of the patients are summarized in Table 1.

### Isolation of plasma and PBMC from human, and RNA extraction

PBMC were prepared using BD Vacutainer^®^ CPT^TPM^ Mononuclear Cell Preparation Tube – Sodium Citrate (BD, Franklin Lakes, NJ, USA) following the manufacturer’s protocol. The obtained samples were centrifuged at 1700 x *g* for 20 min at room temperature within 1 h after blood collection to avoid unwanted deterioration of the samples. The plasma layer was collected and immediately frozen in liquid nitrogen and stored at −80°C until use. The interphase layer (enriched with PBMC) was transferred to a centrifuge tube. D-PBS(-) without Ca and Mg, liquid (PBS) (nacalai tesque, Kyoto, Japan) was added to a final volume of 3 mL, and the tube was centrifuged at 300 x *g* for 10 min at room temperature. The obtained PBMC pellets were washed with 10 mL PBS, lysed in buffer RLT (QIAGEN, Venlo, Netherlands) and stored at −80°C until use. Total RNA was extracted using the RNeasy Mini Kit (QIAGEN) following the manufacturer’s protocol.

### Plasma albumin measurement for human samples

Plasma was collected from human blood samples as described above. Plasma albumin was measured at ORIENTAL YEAST CO., LTD, Tokyo, Japan.

### 4T1 murine breast cancer model

All animal experiment protocols were approved by the Animal Care and Use Committee of Kyoto University. Mice were reared in a 12-h light/dark paradigm with food (CE-2, CLEA Japan, Inc, Tokyo, Japan (https://www.clea-japan.com/products/general_diet/item_d0030) and water available *ad libitum*. Mice were randomly assigned to different experimental groups without any pre-determined criterion. We did not perform any blinding. WT BALB/c mice were purchased from Japan SLC Inc. (Hamamatsu, Japan). 4T1 mouse breast cancer cell line^25^ was cultured and maintained in RPMI1640 (nacalai tesque) supplemented with 10% fetal bovine serum (FBS) (Sigma-Aldrich, Saint Louis, Missouri, USA), 1% penicillin/streptomycin (nacalai tesque) in a 5% CO_2_ tissue culture incubator at 37°C. The thawed cells were passaged once and then were transplanted into mice. 2.5 x 10^6^ 4T1 cells resuspended in 100 μL of RPMI1640 media that were FBS-and penicillin/streptomycin-free were inoculated subcutaneously into the right flank of 8-week-old BALB/c females. In the sham-operated group, mice were given the same amount of RPMI1640 containing neither FBS nor penicillin/streptomycin. Mice were sacrificed 14 days post-transplantation.

### PBMC isolation and RNA extraction from mice

Whole blood samples were collected from the hearts using heparin-coated syringes (heparin-Na, 1000 U/ml; nacalai tesque). PBMC were isolated from mouse whole blood by gradient centrifugation using Histopaque 1083 (Sigma-Aldrich) following the manufacturer’s protocol^57^. Collected whole blood was diluted with an equal volume of PBS, loaded into a 15 mL tube containing 3 mL Histopaque 1083, and centrifuged at 400 x *g* for 15 min at room temperature. The interphase layer (PMBC located between plasma and Histopaque 1083) was collected and washed with 5 mL PBS. The PBMC pellets were immediately lysed in buffer RLT (QIAGEN) or in Trizol reagents, according to the total number of obtained cells, and stored at −80°C until use. Total RNA was extracted using the RNeasy Micro Kit (QIAGEN) or the RNeasy Mini Kit (QIAGEN) following the manufacturer’s protocol.

### Plasma albumin measurement for mice

Whole blood samples were collected from the hearts using heparin-coated syringes (heparin-Na, 1000 U/ml; nacalai tesque) and centrifuged at 1500 x *g* for 20 min at 4°C. The obtained supernatant (plasma) samples were stored at −80°C and plasma albumin was measured at ORIENTAL YEAST CO., LTD, Tokyo, Japan.

### PBMC transcriptome

Total RNA was treated with RNase-free DNase set (QIAGEN). RNA-seq libraries were generated using the NEBNext Globin&rRNA depletion kit and the NEBNext UltraII Directional RNA Library prep kit according to the manufacturer’s protocols (New England Biolabs, MA, USA), as described previously^58^. Sequencing experiments were performed with NextSeq 500 (Illumina; High Output Kit v2.5, 75 Cycles). The obtained reads from human PBMC were mapped to the human reference genome h38/GRCh38 and those from mouse PBMC were mapped to the mouse reference genome mm10/GRCm38 using Hisat2^59, 60^ and counted using Feature Count^61^ in Galaxy (https://usegalaxy.org/). Read counts were normalized using the transcripts per kilobase million (TPM) method, and the genes with expression of 0 and non-protein-coding genes were removed. Gene expression matrices with the obtained TPM scores are listed in Extended Data Table 1 for humans and in Extended Data Table 2 for mice.

### Chart information

Serum albumin levels, C-reactive protein levels, neutrophil counts, and lymphocyte counts of the metastatic breast cancer patients were obtained from the patients’ medical records at Kyoto University Hospital.

### Data analyses

Weighted gene co-expression network analysis (WGCNA): genes with TPM exhibiting more than 1.5-fold changes between metastatic breast cancer patients and healthy volunteers were analyzed using WGCNA. One of the main parameters to be specified by the user in WGCNA is the soft threshold parameter *β*^24^. Among different values of *β* tested, the value *β* = 8 was chosen because it resulted in the best balance between scale-free topology model fit and mean connectivity (Extended Data Fig. 2a, b). The WGCNA function blockwiseModules was used for constructing an unsigned topological overlap matrix to represent the connections between the nodes of the graph with the parameters set as follows: TOMType = “unsigned,” minModuleSize = 30, reassignThreshold = 0, mergeCutHeight = 0.25, numeric labels = TRUE, pamRespectsDendro = FALSE. Pearson correlation coefficients between the module eigengenes and clinical data (albumin scores and age) were then calculated, and the corresponding *p*-values were calculated using Student’s *t*-test. Albumin-correlated modules were defined as those with correlation coefficient |*r*| > 0.5 and *p* < 0.05 with albumin scores. In these modules, correlation coefficients between individual genes and clinical data (albumin scores and age) were calculated using log_2_ values normalized to the average of five healthy volunteers. Module-2 genes with log_2_ normalized scores exhibiting correlation coefficients |*r*| > 0.5 with albumin scores and |*r*| ≤ 0.5 with age were further considered albumin-associated but not age-associated genes.

ImmuCellAI: ImmuCellAI and ImmuCellAI-mouse^30, 31^, bulk RNA-seq data deconvolution approaches, were applied to estimate the abundance of immune cell types in human and mouse PBMC, respectively, using the default settings.

Volcano plots: volcano plots were generated using ggplot2 to visualize differentially expressed genes and metabolites (https://ggplot2.tidyverse.org/index.html). The differentially expressed genes and metabolites were highlighted based on thresholds |log_2_(fold change)| > 0.585 and *p* < 0.05.

### scRNA-seq data analyses from healthy volunteers

Single-cell RNA-seq data were retrieved from the previous publication^26^. Datasets from three Japanese females (*n* = 3) were used for our analysis. Cell annotations, UMAP, and feature plots were performed as described previously^26^.

### Plasma metabolome

Human and mouse plasma samples were prepared for metabolite extraction using the Bligh and Dyer’s method^62^ with minor modifications^13^. Briefly, the metabolites were extracted from plasma (50 μL) with 950 μL of cold methanol (–30°C) containing 10-camphor sulfonic acid (10-camp) (1.25 nmol) and piperazine-1,4-bis (2-ethane sulfonic acid) (PIPES) (1.25 nmol) as internal standards (ISs). The samples were vortexed for 1 min and sonicated for 5 min to ensure thorough mixing, followed by incubation on ice for 5 min to precipitate protein. After centrifugation at 16000 x g for 5 min at 4°C, the supernatant was collected. Then, 600 μL of supernatant was mixed with an equivalent volume of chloroform and 480 μL of water, and then centrifuged at 16000 × g for 5 min at 4°C. The aqueous (upper) layer (720 μL) was transferred to a clean tube for analysis by an ion chromatography (Dionex ICS-5000+ HPIC system, Thermo Fisher Scientific, MA, USA) with a Dionex IonPac AS11-HC-4 μm column (2 mm i.d. × 250 mm, 4 μm particle size, Thermo Fisher Scientific) coupled with a Q Exactive, high-performance benchtop quadrupole Orbitrap high-resolution tandem mass spectrometer (Thermo Fisher Scientific) (IC/MS) for anionic polar metabolites (i.e., organic acids, nucleotides, etc.)^63^, a liquid chromatography (Nexera X2 UHPLC system, Shimadzu Co., Kyoto, Japan) with a Discovery HS F5 column (2.1 mm i.d. × 150 mm, 3 μm particle size, Merck, Darmstadt, Germany) coupled with a Q Exactive instrument (PFPP-LC/MS) for cationic polar metabolites (i.e. amino acids, bases, nucleosides, etc.)^63^, or a LC (Shimadzu Co.) with a GL-HilicAex column (2.1 mm i.d. × 150 mm, 5 μm particle size, Resonac Techno Service Co., Ibaraki, Japan) coupled with a Q Exactive instrument (unified-hydrophilic-interaction/anion-exchange liquid chromatography mass spectrometry, unified-HILIC/AEX/MS) for cationic and anionic polar metabolites^64^. The aqueous layer extracts were evaporated under vacuum and stored at −80°C until used for unified-HILIC/AEX/MS analysis of human plasma extracts and IC/MS and PFPP-LC/MS analyses of mouse plasma extracts. Prior to analysis, the dried aqueous layer was reconstituted in 50 μL of water. LC/MS data analysis was performed using Multi-ChromatoAnalysT (BeForce Co., Fukuoka, Japan) and Cascade (Reifycs Inc., Tokyo, Japan).

Expression of the hydrophilic metabolites was calculated using peak area relative to the internal standards of the HRMS precursor 10-camp as described previously^58^. In Fig. 6a, b, we also used scores normalized to PIPES to select metabolites whose correlations to albumin (|*r*| > 0.5 and *p* < 0.05) were consistent between 10-camp and PIPES. Expression of metabolites are listed in Extended Data Table 3 for humans and in Extended Data Table 4 for mice.

### Statistical analysis and data visualization

The sample size was determined based on feasibility. GraphPad Prism Software (Prism9) was used to analyze data. Data are displayed as mean ± SEM. Student’s *t*-test was performed to analyze the statistical significance and *p* < 0.05 was considered statistically significant. Simple regression analysis was performed to obtain the Pearson correlation coefficients and *p*-values. Graphs were generated using GraphPad Prism Software and R. Cartoons in Fig. 1a and Fig. 6g were obtained from BioRender (https://www.biorender.com).

## Data availability

The bulk RNA transcriptome datasets used in this study are available in DNA Databank of Japan (DDBJ) under the accession numbers of DRA016869 (human PBMC transcriptome) and DRA016870 (mouse PBMC transcriptome). scRNA-seq datasets retrieved from the previous publication^26^ are available at https://ddbj.nig.ac.jp/resource/jga-study/JGAS000321.

## Acknowledgement

This work was supported by JSPS KAKENHI (20H03451; S.K.: 19K16770 and 21K15530; K.K.), JST FOREST (20351876; S.K), AMED (JP21ck0106698; K.K.), JST Moonshot (JPMJMS2011-61; S.K), Caravel, Co., Ltd (S.K), and Japan Foundation for applied Enzymology (SK). This work was also partly performed in the Medical Research Center Initiative for High Depth Omics Program of the Medical Institute of Bioregulation (MIB) and the Cooperative Research Project Program of the MIB, Kyushu University.

We thank M. Goto and T. Nakaji (Kyushu University) for their technical supports on this work.

## Competing interests

KK: grants from TERUMO, Astellas, Eli Lilly, Kyoto Breast Cancer Research Network; consulting fee from Becton Dickinson Japan; honoraria from Eisai, Chugai, and Takeda MT: grants from Chugai, Takeda, Pfizer, Taiho, JBCRG Assoc., KBCRN Assoc., Eisai, Eli Lilly, Daiichi-Sankyo, AstraZeneca, Astellas, Shimadzu, Yakult, Nippon Kayaku, AFI Technology, Luxonus, Shionogi, GL Science, and Sanwa Shurui; honoraria from Chugai, Takeda, Pfizer, Kyowa-Kirin, Taiho, Eisai, Daiichi-Sankyo, AstraZeneca, Eli Lilly, MSD, Exact Science, Novartis, Shimadzu, Yakult, Nippon Kayaku, Devicore Medical Japan, Sysmex; Advisory Board of Daiichi-Sankyo, Eli Lilly, BMS, Athenex Oncology, Bertis, Terumo, Kansai Medical Net; Board of Directors of JBCRG Assoc., KBCRN, NPO org. OOTR, and JBCS Assoc; Associate Editor of the British Journal of Cancer, Scientific Reports, Breast Cancer Research and Treatment, Cancer Science, Frontiers in Women’s Cancer, Asian Journal of Surgery, Asian Journal of Breast Surgery.

All remaining authors declare no conflicts of interest.

## Author contribution

Y.N. collected patient samples, performed mouse experiments, analyzed data, constructed figures, and contributed to the writing of the manuscript. M.Y. performed transcriptome measurements. Y.K. and Y.S. performed scRNA-seq data analyses. Y.I., M.T., K.N., and T.B. performed metabolome measurements. R.K. and C.H. performed mouse experiments. D.P.S. and A.V. supported data analyses. M.T. contributed to conception of this study. K.K. conceived and supervised the study, designed experiments, and collected patient samples. S.K. conceived and supervised the study, designed experiments, analyzed data, constructed figures, and wrote the manuscript. All authors reviewed the manuscript and provided intellectual inputs.

