## Supplementary information for "Low albumin status accompanies multi-layered immunosuppressive phenotypes in metastatic breast cancer patients"

**Extended Data Figure Legends**

**Figure 1: Serum albumin and CRP levels in our patient cohort**

**a.** Correlation between serum albumin levels and plasma albumin levels in metastatic breast cancer patients ($n$ = 10). Note that there are three patients having the same albumin scores (plasma albumin, serum albumin) = (3.6, 3.8).

**b.** Correlation between plasma albumin levels and C-reactive protein (CRP) levels in metastatic breast cancer patients ($n$ = 10). Note that there are three patients having the same scores (albumin, CRP) = (3.6, 0.1).

**a-b.** The Pearson correlation coefficients and $p$-values obtained by simple regression analysis (GraphPad Prism) are shown.

**Figure 2: Weighted gene co-expression network analyses and its characterization**

**a.** Scale independence in the WGCNA analysis.

**b.** Mean connectivity in the WGCNA analysis.

**c.** Volcano plot of $\log_2$ fold average (4T1/sham) versus $-\log_{10}$ ($p$-value) is shown. The volcano plot is generated from PMBC RNA-seq from sham-operated and 4T1-bearing mice 14 days after 4T1 transplantation. Genes showing more than 1.5-fold change and $p < 0.05$ are highlighted. $n = 4$.

**d.** Bar plot showing $\log_2$ fold changes of module-3 genes between sham-operated and 4T1-bearing mice. Genes showing the same direction of changes in the 4T1 breast cancer model compared to the human datasets are shown. Genes upregulated in 4T1-bearing mice are shown in red. Genes downregulated in 4T1-bearing mice are shown in blue. Genes are ordered according to correlations of corresponding human genes to albumin levels in humans. The names of the top10 genes most strongly negatively correlating with albumin levels in humans are indicated.

**Figure 3: Prediction of changes in immune cell abundances in cancer-bearing conditions**

**a.** UMAP plot of scRNA-seq from Japanese healthy volunteers.

**b.** UMAP plot for *S100A8*.

**c.** Estimated abundance of neutrophils in PBMC in the 4T1 breast cancer model calculated by ImmuCell-AI-mouse.

**d.** Estimated abundance of monocytes in PBMC in the 4T1 breast cancer model calculated by ImmuCell-AI-mouse.

**e.** Estimated abundance of human dendritic cells in PBMC calculated by ImmuCell-AI and its correlation to plasma albumin levels.

**f.** Estimated abundance of dendritic cells in PBMC in the 4T1 breast cancer model calculated by ImmuCell-AI-mouse.

**g.** Estimated abundance of human macrophages in PBMC calculated by ImmuCell-AI and its correlation to plasma albumin levels.

**h.** Estimated abundance of human natural killer T cells in PBMC calculated by ImmuCell-AI and its correlation to plasma albumin levels.

**i.** Estimated abundance of human $CD4^+$ T cells in PBMC calculated by ImmuCell-AI and its correlation to plasma albumin levels.

**a-b.** The data are retrieved from previously published single-cell RNA-seq (scRNA-seq) datasets from PBMC of Japanese healthy volunteers ($n$ = 3 (females))[26]. Previously annotated monocyte cluster abundantly express a neutrophil marker *S100A8*, being considered as neutrophils.

**c-d, f.** Data are presented as the mean ± SEM. The $p$-value is shown (non-paired, two-tailed Student $t$-test). $n$ = 4 for sham-operated and 4T1-bearing mice. PBMC was collected 14 days after 4T1 transplantation.

**e, g-i.** The Pearson correlation coefficients and $p$-values obtained by simple regression analysis (GraphPad Prism) are shown.

**Figure 4: Immunosuppressive phenotypes correlate with low albumin levels**

**a.** Estimated abundance of $CD8^+$ T cells in PBMC in the 4T1 breast cancer model calculated by ImmuCell-AI-mouse.

**b.** Estimated abundance of human natural killer cells in PBMC calculated by ImmuCell-AI and its correlation to plasma albumin levels. The Pearson correlation coefficients and $p$-values obtained by simple regression analysis (GraphPad Prism) are shown.

**c.** Estimated abundance of $\gamma\delta$ T cells in PBMC in the 4T1 breast cancer model calculated by ImmuCell-AI-mouse.

**d.** Estimated abundance of natural killer cells in PBMC in the 4T1 breast cancer model calculated by ImmuCell-AI-mouse.

**a, c, d.** Data are presented as the mean ± SEM. The *p*-value is shown (non-paired, two-tailed Student *t*-test). *n* = 4 for sham-operated and 4T1-bearing mice. PBMC was collected 14 days after 4T1 transplantation.

**Figure 5: Neutrophil-to-lymphocyte ratio correlates with low albumin levels**

Correlation between plasma albumin levels and neutrophil-to-lymphocyte ratio calculated by the data retrieved from patients' chart. The Pearson correlation coefficients and *p*-values obtained by simple regression analysis (GraphPad Prism) are shown.

**Figure 6: Characterization of plasma metabolome in cancer-bearing conditions**

**a.** Expression of tryptophan.

**b.** Expression of kynurenine.

**a-b.** Fold change data normalized to the average of healthy volunteers are presented as the mean ± SEM. The *p*-value is shown (non-paired, two-tailed Student *t*-test). *n* = 5 for HV, *n* = 10 for MBC.

**Extended Data Tables**

Extended Data Tables 1-6 are provided as separate excel files.

### Extended Data Figure 1

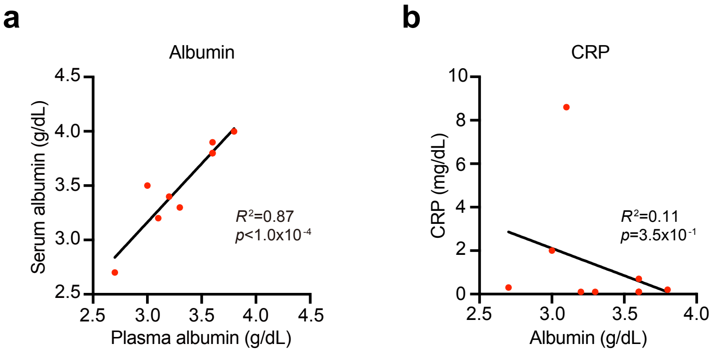

### Extended Data Figure 2

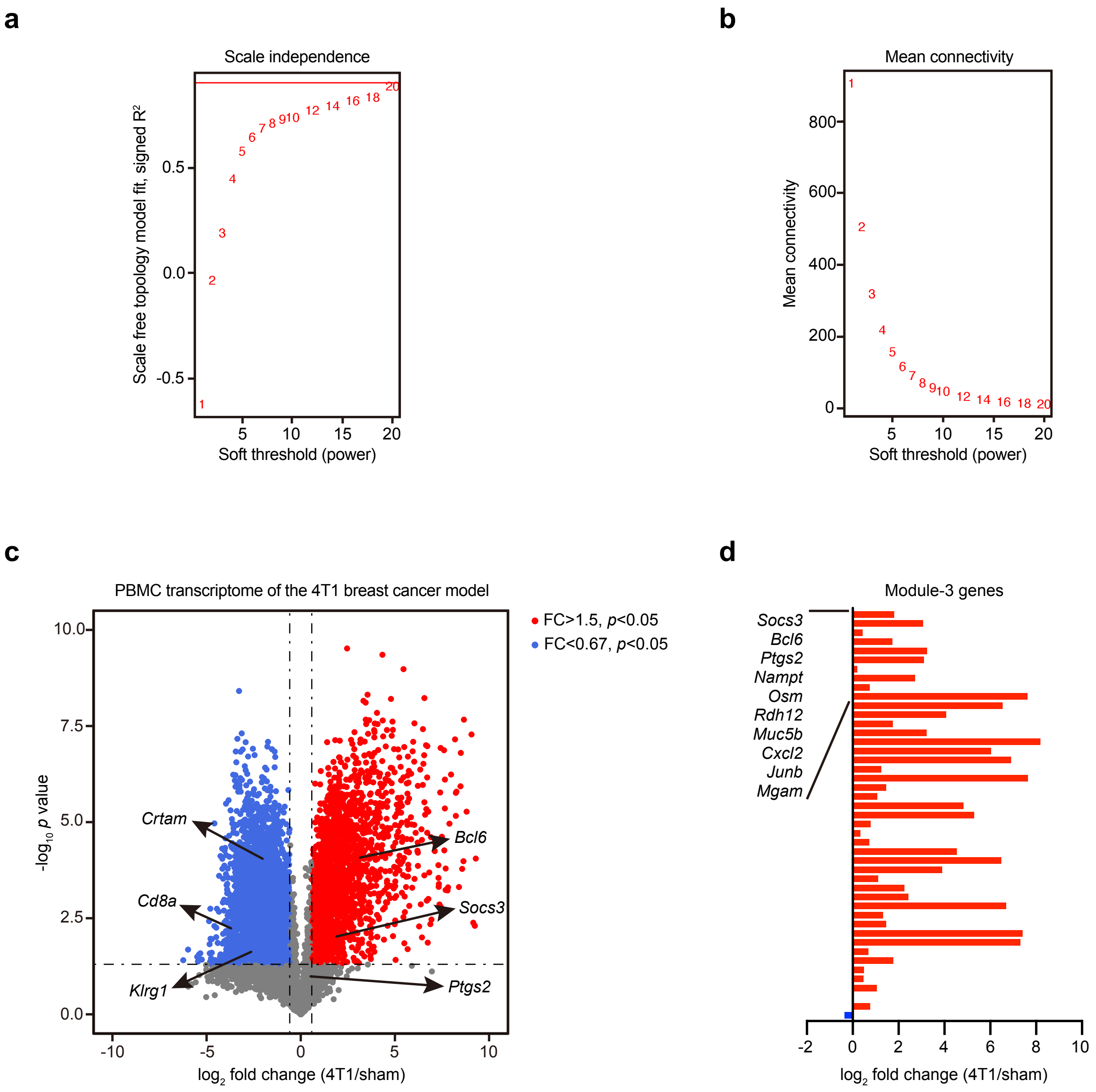

#### Extended Data Figure 3

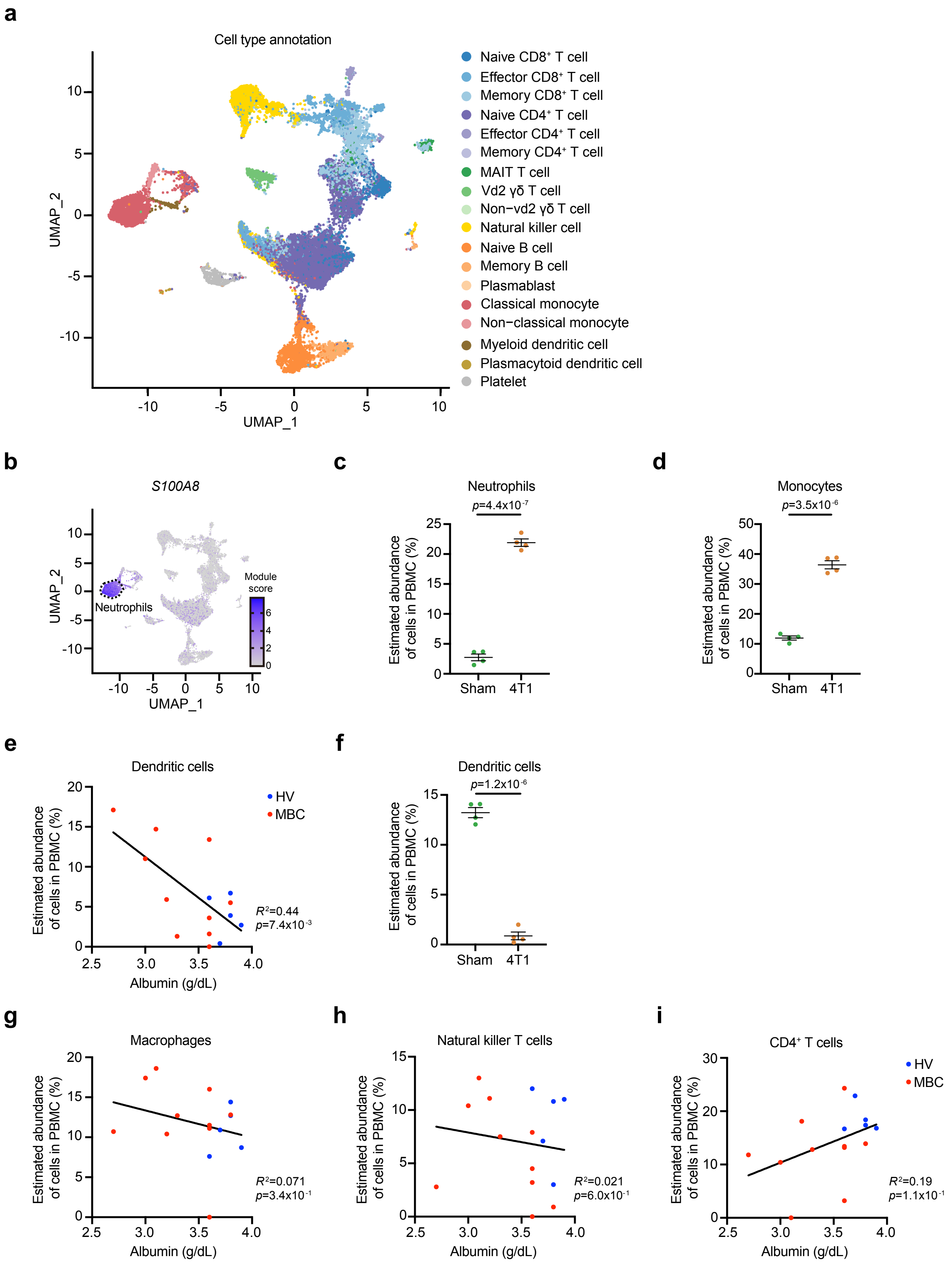

### Extended Data Figure 4

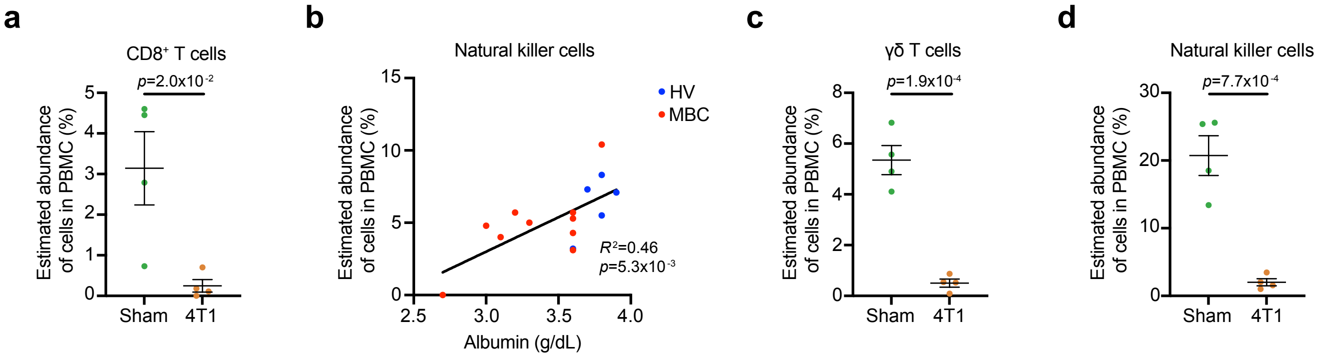

### Extended Data Figure 5

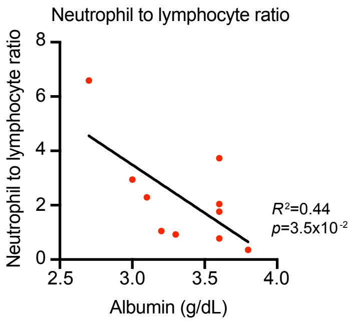

### Extended Data Figure 6

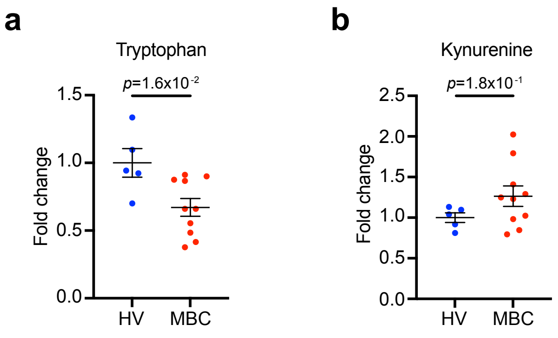
